# The Role of Fetuin-A on the Attachment and Proliferation of Osteoblast-like Cells on Model Gold Surfaces

**DOI:** 10.1101/2025.11.04.686303

**Authors:** Alessandra Merlo, Jesper Medin, Shane Scott, Andreas Dahlin, Kathryn Grandfield, Kyla N. Sask

## Abstract

Fetuin-A is a plasma protein of interest for bone-interfacing applications due to its role in mineralization processes through calcium/phosphate ion-binding capabilities. However, the role of fetuin-A in the initial stages of cellular interaction with biomaterials and the mechanisms involved are not fully clear. This work investigated the response of osteoblast-like Saos-2 cells to model gold substrates presenting pre-adsorbed fetuin-A as a surface modification, to determine the role of the protein in cell attachment and proliferation. Correlative quartz crystal microbalance with dissipation (QCM-D), surface plasmon resonance, and radiolabeling confirmed fetuin-A adsorbed on model surfaces in similar quantities compared to serum albumin but formed a less packed layer with increased water entrapment. Surfaces presenting pre-adsorbed fetuin-A enhanced cellular adhesion, similar to fibronectin, but attached cells displayed morphological characteristics more similar to those with pre-adsorbed albumin, with lower average surface area and maximum axis. Over 3 days, fetuin-A exhibited lower cellular proliferation compared to the fibronectin control, likely correlated to the decrease in cellular metabolism observed at the same time-point, and persisted over 7 days. These results provide insight into the role of adsorbed fetuin-A for bone-interfacing implant applications, suggesting the pre-adsorption of the protein alone aids cellular attachment, but is not sufficient to promote early stages of osseointegration.

## 1. Introduction

The need for bone-interfacing implants for orthopedic and dental applications is increasing due to the aging population, yet there are still significant issues with these, such as bacterial colonization^1^, bone resorption, and lack of, or poor, osseointegration ^1,2^. To overcome these problems, a variety of methods have been investigated, based on the established concept that surface characteristics of the implant materials impact the first stages of the body’s response ^3–5^, both in regulating the inflammatory response ^6–8^ and the subsequent osteogenic and angiogenic processes ^9,10^. Among the most common methods used to promote osseointegration are surface modifications aimed at varying either or both the chemistry and the topography of the surface, combining various methods such as sand blasting, chemical etching, oxidative processes and laser treatment ^11–13^. Alternatively, the deposition of specific biomolecules^14–16^ or biomimetic^17^ coatings, such as hydroxyapatite^18,19^, polydopamine^10^ or specific moieties^20^, can provide improved cell adhesion and proliferation and promote tissue regeneration.

However, one aspect often overlooked in osseointegration is that, upon insertion of the implant inside the body, the biomaterial does not initially contact the cells or tissue. Instead, upon implantation, the surface will first come in contact with biological fluids, such as blood, and be coated by an adsorbed layer of plasma or other proteins^21^, which mediate the following stages of the integrative process, including inflammatory response, bone cell recruitment and attachment, proliferation, and differentiation, as well as osteogenic and angiogenic processes ^22^. As such, it is the adsorbed layer of proteins, its composition and spatial arrangement, that determine the subsequent biological response and define the first stages in the osseointegrative process.

Fetuin-A, also named α2-Heremans-Schmid glycoprotein (AHSG), is a blood plasma glycoprotein measuring around 60 kDa with numerous physiological roles^23^. It has been reported to be involved in the insulin signaling pathway^24,25^ and incidence of type 2 diabetes^26^, to act as either a pro-or anti-inflammatory agent in different immunoresponses^27,28^, as well as in aiding fetal brain development^29^. Among its various functions, however, fetuin-A has elicited interest in current bone-interfacing applications, given its participation in mineralization processes and bone metabolism^30–32^. Specifically, fetuin-A is pivotal for both intrafibrillar and extrafibrillar mineralization by acting as a mineralization inhibitor^33^, and serves as both a scavenger and carrier of calcium and phosphate ions by forming protein-mineral complexes known as calciprotein particles (CPPs)^34^.

While the role of Fetuin-A and its interaction with the inorganic component of bone have been more thoroughly investigated^23,31,35^, the physiological role of fetuin-A in connection with the organic components of bone, and particularly its role on bone cells response, still needs to be explored. Oschatz et al^36^ reported an increase in cell attachment and spreading of osteoblast-like cells when fetuin-A was pre-adsorbed on nonwoven PLLA-co-PEG electrospun mats. In addition, the presence of fetuin-A allowed for higher hydroxyapatite-precursor deposition from supersaturated calcium phosphate solution, but no difference was highlighted in cellular metabolic activity, nor in collagen-I synthesis, with respect to the non-preadsorbed surfaces. Most other previous investigations have assessed fetuin-A in blood, given its presence in plasma, leaving limited reports in relation to its adsorption at the biointerface and resulting effects on cells. Therefore, further studies are needed to assess the direct role of fetuin-A on attachment and proliferation of osteoblast cells to other materials, and to determine the protein characteristics when the physicochemical and mechanical properties of the surface are varied, for example in implants that are commercially available.

The present investigation compared the quantity of protein adsorbed, kinetics of adsorption, viscoelastic properties of the adsorbed layer of fetuin-A on model gold substrates using correlative quartz-crystal microbalance (QCM), surface plasmon resonance (SPR), fluorescence microscopy imaging, and radiolabeling, to address homogeneity of the protein layer and gain thorough insight into the adsorption processes of this protein. Fetuin-A was compared to serum albumin, similar in both molecular weight and structure in solution^36,37^.The response of the osteoblast-like cell line Saos2 to pre-adsorbed fetuin-A was compared to pre-adsorbed albumin (negative control) and fibronectin (positive control). Pre-adsorbed substrates were analyzed for cellular attachment, cell morphology, proliferation and metabolic activity over 7 days, to address any inherent capability of fetuin-A in promoting earlier start of the osseointegrative process.

## 2. Materials and Methods

### 2.1 Materials

Bovine fibronectin (BFn) was purchased from Sigma-Aldrich (F4759) and reconstituted in sterile water (Wisent Inc., St. Bruno, QC, CA) as per the manufacturer’s directions. Bovine serum albumin (BSA, 05470) and bovine fetuin (BFet, F2379) were purchased from Sigma Aldrich and dissolved in phosphate buffered saline (PBS) (Wisent Inc) in stock solutions of 1 mg/mL. All protein solutions were diluted to a final concentration of 0.2 mg/mL using PBS. McCoy Media 5A (M9309), penicillin/streptomycin (P4333), saponin (S4521) and formaldehyde 37% solution (F8775) were purchased from Sigma-Aldrich (Oakville, ON, CA). For fluorescent protein labelling, Alexa 488 NHS ester (A20000, Thermo Fisher Sci.) fluorescent dye was dissolved at a concentration of 10 mg/mL in dimethyl sulfoxide (DMSO), while BSA and BFet were dissolved at concentrations of 6 mg/mL in 0.2 M NaHCO_3_ (S8875, Sigma-Aldrich) at pH 8.3.

### 2.2 Surface Preparation and Modification

Gold substrates composed of 525±25 µm-thick silicon wafers were coated with a 300-500 Å chromium adhesive layer followed by a layer of gold (approximately 1000 Å) then diced into 0.5 x 0.5 cm pieces (Silicon Valley Microelectronics Inc, Buellton, CA). The gold-coated silicon was cleaned by immersion in a boiling solution of Milli-Q water, hydrogen peroxide, and ammonium hydroxide (4:1:1 v/v) for 10 min followed by rinsing in Milli-Q water and drying under nitrogen. To adsorb proteins to gold, the surfaces were incubated in 0.2 mg/mL protein solutions dissolved in PBS at RT under static conditions for 3 h.

### 2.3 Water Contact Angle Measurements

For water contact angle measurements, substrates produced as described in Section 3.2, but diced in 1 x 1 cm pieces, were used. Water contact angle measurements were performed using the sessile drop method with a Ramé-Hart NRL 100-00 goniometer (Mountain Lakes, NJ, USA). Data are reported as average ± SD, with N≥12 (where N is the number of replicates), and analyzed using the one-way Analysis of Variance (ANOVA) method. The significance level was defined at 0.05, with *p<0.05, **p<0.01, and ***p<0.001.

### 2.4 Quartz-Crystal Microbalance with Dissipation

Quartz Crystal Microbalance with Dissipation (QCM-D) gold sensors (QuartPro, Stockholm, Sweden; 5MHz Cr/Au) were cleaned as described in Section 2.2. Measurements were performed on a QSense Analyzer (Biolin Scientific, Gothenburg, Sweden) in PBS buffer (pH 7.4) following baseline stabilization for at least 20 min. The flow rate of buffer used was 100 μL/min, to conduct the observation under low flow conditions, similar to the static protein immobilization used in Section 2.2, and the measurements were done at 25°C. The mass adsorbed on the sensor was estimated through the Sauerbrey equation (Equation 1) using the fifth overtone and a *C* value of 17.7 ng/(cm^2^*Hz).

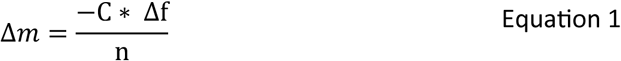

The association constants for both proteins were obtained using the Langmuir adsorption model (Equation 2): with the assumption C(t) is constant at C_0_ and that (t) ≈ 0 for small t values, the association constant can be calculated (as described in the 1S.1 section of the Supplemental Information) considering the value of 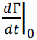 equal to the initial slope of the SPR in-situ real-time curves. All modelled values are provided with a 95% confidence interval.

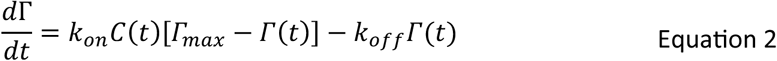

The association constants for both proteins were obtained using initial slope values. The total protein adsorbed mass reported, as well as the association constant, refer to the calculations from the 5^th^ overtone with maximum shift measured after 40 min, divided by the overtone number, with N≥2 replicates. To confirm the total amount adsorbed after 3 h adsorption and full surface coverage, longer runs of 160 min were conducted with the same parameters.

### 2.5 Surface Plasmon Resonance

SPR sensors (BioNavis, Tampere, Finland) were cleaned as described in Section 2.2. Measurements were performed on a SPR Navi 220A instrument (BioNavis) in PBS buffer (pH 7.4) following baseline stabilization for at least 20 min. The total internal reflection (TIR) and SPR angle were monitored with a 670 nm laser diode. The flow rate of buffer used was 10 μL/min, to conduct the observation under low flow conditions, similar to the static protein immobilization used Section 2.2, and the measurements were done at 25°C. The total *in situ* real-time protein adsorbed mass reported refers to the calculations on the 670 nm laser channel done according to previous work^39^, with maximum shift measured after 40 min (N≥2 replicates). For correlative estimates, the total mass of adsorbed protein after 3 h of protein immobilization under static conditions was confirmed with measurements in dry conditions (N≥2) with the 670 nm laser diode, followed by rinsing in PBS and drying in nitrogen. SPR spectra analysis on dry measurements by Fresnel modeling was performed according to the methodology previously reported^39^.

### 2.6 Protein Radiolabelling with ^125^Iodine

BSA and BFet were labelled with Na^125^I using the iodine monochloride (ICl) method. The labelled protein was filtered with an AG 1-X4 anion exchange resin to eliminate free ^125^I. Labeled protein was precipitated in 20% trichloroacetic acid (TCA) and centrifuged for 5 min. The supernatant, containing free iodide ion (i.e. not bound to protein) was counted on a Wizard Automatic Gamma Counter (PerkinElmer, Shelton, CT, USA). Tests were conducted to determine residual free iodine, and levels were maintained below 3%. To prevent the binding of residual free iodine to the gold substrate, protein dilution and adsorption experiments were carried out using phosphate buffered saline containing nonradioactive sodium iodide (PBS–NaI) (3%) ^40^, with labelled and unlabelled protein in a 1:9 mass ratio. BSA and BFet solutions in PBS-NaI buffer were prepared with final concentrations of 0.2 mg/mL, and samples were incubated for 3 h at room temperature in static conditions. Following adsorption, the samples were rinsed three times with PBS, for 5 min each. The samples were lightly wiped to remove the residual adherent buffer, and the radioactivity was counted with a gamma counter. Radioactivity was converted to protein adsorption amounts based on corresponding solution counts. 4 replicates for each condition were used. Following radioactivity counting, samples were incubated in sodium dodecyl sulfate (SDS, 2% in water) overnight under shaking conditions (250 rpm) to elute loosely adsorbed protein. The SDS solution was subsequently removed, and the sample surfaces were rinsed three times with PBS buffer for 5 min each. Finally, the samples were dried, and radioactivity measured through gamma radiation counts as previously described to obtain the residual surface coverage.

### 2.7 Fluorescently-tagged Protein Surface Quantification

To covalently bind the Alexa488 fluorophore to proteins, the protein and Alexa488 NHS solutions were mixed in a ratio of 50:1 and left to react for 1 h while protected from light. Solutions containing the Alexa488-labelled proteins and free dye were purified from unreacted dye by pipetting them onto Zeba spin-dye columns (A44296, Thermo Fisher Sci.) and centrifuging for 2 min at 1,000xg, with the solutions containing the fluorescently-tagged proteins contained in the elute. A description of protein concentration quantification and the degree of labelling determination and calculation is provided in SI section 1S.2. Fluorescent BSA and BFet were deposited on gold substrates as described in Section 2.2 and protected from light. Samples were washed in PBS 3 times for 5 min, then dried using nitrogen. A glass slide was cleaned using soapy water and a scrub brush, dried using nitrogen, and samples were placed face down on the slide. Fluorescence microscopy was performed using a 20x microscope objective equipped on a Cytation 7 (Agilent) multimode reader using the inverted microscopy setting, and imaging with the GFP cube (469/35 nm excitation, 525/39 nm emission) at 5 Intensity, 200 ms integration time, and 30 Gain. Three different images on each sample were collected. Image analysis was performed using Fiji^41^ and calculations using Matlab software. The normalized average fluorescence intensity <I_protein, norm_> and average coefficient of variation <CV> for each protein-functionalized gold sample were calculated as described in SI Section 1S.2.

### 2.8 Cell Culture

Saos-2 osteosarcoma cells (ATCC®, HTB-85) were grown in McCoy’s 5A modified medium with 15% fetal bovine serum (FBS, Wisent Inc., St. Bruno, QC, CA) and 1% penicillin/streptomycin. All experiments were conducted with cell passage number ≤10. All samples were cleaned as described in section 3.2 and then sterilized with 30 min exposure to UV light on both sides of the substrates. To seed cells on the substrates, cells were treated with 0.25% Trypsin-EDTA for 6 min at 37°C. Immediately afterwards, an equal amount of media was added and the cells were centrifuged for 5 min at 1000xg, the eluted solution removed, and the cell pellet resuspended in non-supplemented media. Cells were seeded in non-supplemented media and incubated for 24 h at 37 °C with 5% CO_2_. After 24 h, non-supplemented media was substituted for McCoy’s 5A modified medium supplemented with 15% FBS and 1% PS, and culture media exchanged every two days.

### 2.9 Fluorescence Microscopy

Cell attachment and proliferation were assessed using immunostaining on days 1, 3, and 7. Before staining, substrates were washed twice with PBS to eliminate non-adherent cells present on the surface. Next, substrates were flash permeabilized with a 0.1% solution of saponin for around 10-15 sec, and fixed with a 4% formaldehyde solution for 7 min at 4°C. Samples were washed twice with PBS. A 2% BSA blocking solution was added for 1 h at 37°C. Substrates were washed twice with PBS, and a staining solution containing nucleic stain (Hoechst 33342, Life Technologies Inc., CA, USA) and phalloidin stain (phalloidin-Alexa fluorophore 488) in PBS (3:1:3000 v/v) was added for 2 h at room temperature. Samples were washed with PBS and then mounted face-down with Fluoromount-G Mounting Medium (00-4958-02, Life Technologies Inc., CA, USA) on glass coverslips. Cells were imaged on a Nikon A1R Inverted Confocal with a 20x air objective lens (NA 0.75) on day 1, using the large image function embedded in the NIS Viewer software to image a larger surface area. On day 3 and day 7, a 10x air objective lens, (NA 0.45) was used.

### 2.10 Image Analysis

#### 2.10.1 Cell Nuclei Count

For the cell nuclei count on days 1 and 3, the free software CellProfiler was employed, with the following workflow: the DAPI channel of collected images was thresholded in 2 classes using the Ostu method, and nuclei identified through the Identify Primary Objects functions using the same thresholding method discarding objects <30 and >350 px outside of the size range of single nuclei. To normalize all data over the same surface area, for day 1, stitched images of 20x captures obtained through the “Large Image” option on the Nikon software were cropped to display equal surface area of the 10x images. Day 3 images were used as collected and analyzed with CellProfiler with the same methodology described above. For each condition, ≥3 images on random areas of 3 replicates, per condition, were averaged, and then compared with other substrates. For day 7 images, the confluency over the monolayer required a first segmentation stage in Ilastik to separate nuclei from background through a Pixel Classification process. Output masks were then opened in Fiji software, thresholded with the Ostu method to create a binary mask and then the number of objects counted through the Analyze Particles function with the same thresholds mentioned above. For each condition, ≥3 images on random areas of 3 replicates, per condition, were averaged, and then compared with other substrates. Statistical differences were measured using an independent two-way ANOVA with Tukey’s HSD post-hoc test in GraphPad Software (San Diego, CA, USA). The significance level was defined at 0.05.

#### 2.10.2 Single Cell Surface Area, Eccentricity and Major Axis (Day 1)

To better assess any role played by the protein upon cellular attachment, images at 20x collected on day 1 were employed to evaluate average surface area, eccentricity, major and minor axis of single cells. To the CellProfiler workflow described in Section 2.9.1, an additional module was added to threshold the single cells on the merged channels images: once nuclei were identified through the Identify Primary Object function, the Identify Secondary Objects function was used starting from the identified nuclei as input, using Propagation as identification method and Minimum Cross-Entropy as thresholding method. All images/analysis output produced by the workflow were then checked individually to exclude erroneous thresholds by the software, such as two cells merged into one, or one cell considered two different entities. Only single-standing, isolated cells, with their area fully delimited within the borders of the image, were considered. The revised results were then pooled by condition to consider n≥100 cells per condition. Statistical differences were measured using a one-way ANOVA with Tukey’s HSD post-hoc test in GraphPad Software (San Diego, CA, USA). The significance level was defined at 0.05. Statistical differences were measured using a one-way ANOVA with Tukey’s HSD post-hoc test in GraphPad Software (San Diego, CA, USA). The significance level was defined at 0.05.

#### 2.10.3 Cell Surface Coverage

The Ilastik software was used to evaluate the total surface coverage (% of total area) of the cells using the Pixel Classification method. 3 random images of the dataset, FITC channel, were used to define two classes, cell area and background, and train the model, which was then applied to the full dataset. Exported segmentations were then loaded in Fiji, images converted into binary masks, and pixel for the two categories counted to define surface coverage. For day 1, stitched images of 20x captures obtained through the “Large Image” option on the Nikon software were cropped to display equal surface area of the 10x images. 3 regions of interest per sample were averaged and compared with the averages of other conditions. For day 3 and day 7, images collected were used as is for the Ilastik segmentation, and for each condition, ≥3 random regions of interest of the single sample were averaged, with 3 replicates per sample, and then statistically compared with other substrates. Results are presented as average ± SD. Statistical differences were measured using a two-way independent ANOVA with Tukey’s HSD post-hoc test in GraphPad Software (San Diego, CA, USA). The significance level was defined at 0.05.

### 2.11 Cell Metabolism

Cell metabolism was assessed with an alamarBlue^TM^ assay (Life Technologies Inc., Carlsbad, CA, USA) on days 1, 3 and 7, with 6 replicates for each condition and two repeats. An Infinite M200Pro plate reader (Tecan Group Ltd., Switzerland) was used to measure the fluorescence intensity values at 540 nm excitation and 580 nm emission wavelengths. The values of the two repeats were normalized over the average of day 1 – control value (polystyrene plate) of the corresponding run, then pooled together. Statistical differences were measured using a two-way independent ANOVA with Tukey’s HSD post-hoc test in GraphPad Software (San Diego, CA, USA). The significance level was defined at 0.05.

## 3. Results and Discussion

The role of fetuin-A on bone mineralization has been widely investigated. Given its high number of sialic residues, fetuin-A presents various highly negatively-charged regions capable of binding Ca^2+^ and calcium phosphate ions, forming protein-inorganic CPPs^42–44^. Through this mechanism, fetuin-A allows the storing of calcium and phosphate ions, needed for mineralization processes, by reducing their concentration in body fluids, thus preventing ectopic calcification of bone^45^. The role of fetuin-A on the organic component of bone, and on cellular attachment and proliferation of bone cells is a subject yet to be thoroughly investigated, particularly when fetuin-A is adsorbed on surfaces compared to its presence in biological fluids. A previous study by Oschatz et al used fetuin-A to functionalize nonwoven electrospun PLLA-co-PEG mats^36^. The results of the study suggest pre-bound fetuin-A participates in the regulation of cellular behavior, improving cell attachment and proliferation over 3 days, while no change was observed in cellular metabolism and collagen I synthesis^36^.

In the present work, we aimed to investigate the effect of pre-adsorbed fetuin-A on the attachment and proliferation of the osteosarcoma cell-line Saos2, representative of osteoblast-like behavior^46^ on model gold surfaces. The resulting cellular response is compared to that of the same model gold substrates with either pre-adsorbed albumin, or pre-adsorbed fibronectin. Albumin, the most abundant blood plasma protein, is a commonly used protein for passivating layers, and here serves as a negative control. Fibronectin, is a plasma protein with known cell-adhesive capabilities, and serves as a positive control^47–49^.

This work aims to better elucidate the role of fetuin-A on the attachment and proliferation of osteoblast-like cells. To consider possible differences in structural arrangement, which could impact the exposure of amino acid sequences depending on the binding to the substrate, the investigation compares model gold surfaces, where proteins are attached through adsorption phenomena. The use of gold substrates allows a variety of characterization techniques to be used to obtain correlative information.

### Substrate Characterization

Sessile drop water contact angle (WCA) measurements were performed to assess the protein immobilization and homogeneity of the protein coverage. The results for the WCA analysis are presented in Figure 1. No statistically significant differences were found between the bare gold (gold) and the surfaces with pre-adsorbed albumin (gold+BSA) and fetuin (gold+BFet). A significant difference however was detected between gold and gold with pre-adsorbed fibronectin (gold+BFn), where the contact angle increased by about 15°.

**Figure 1.**
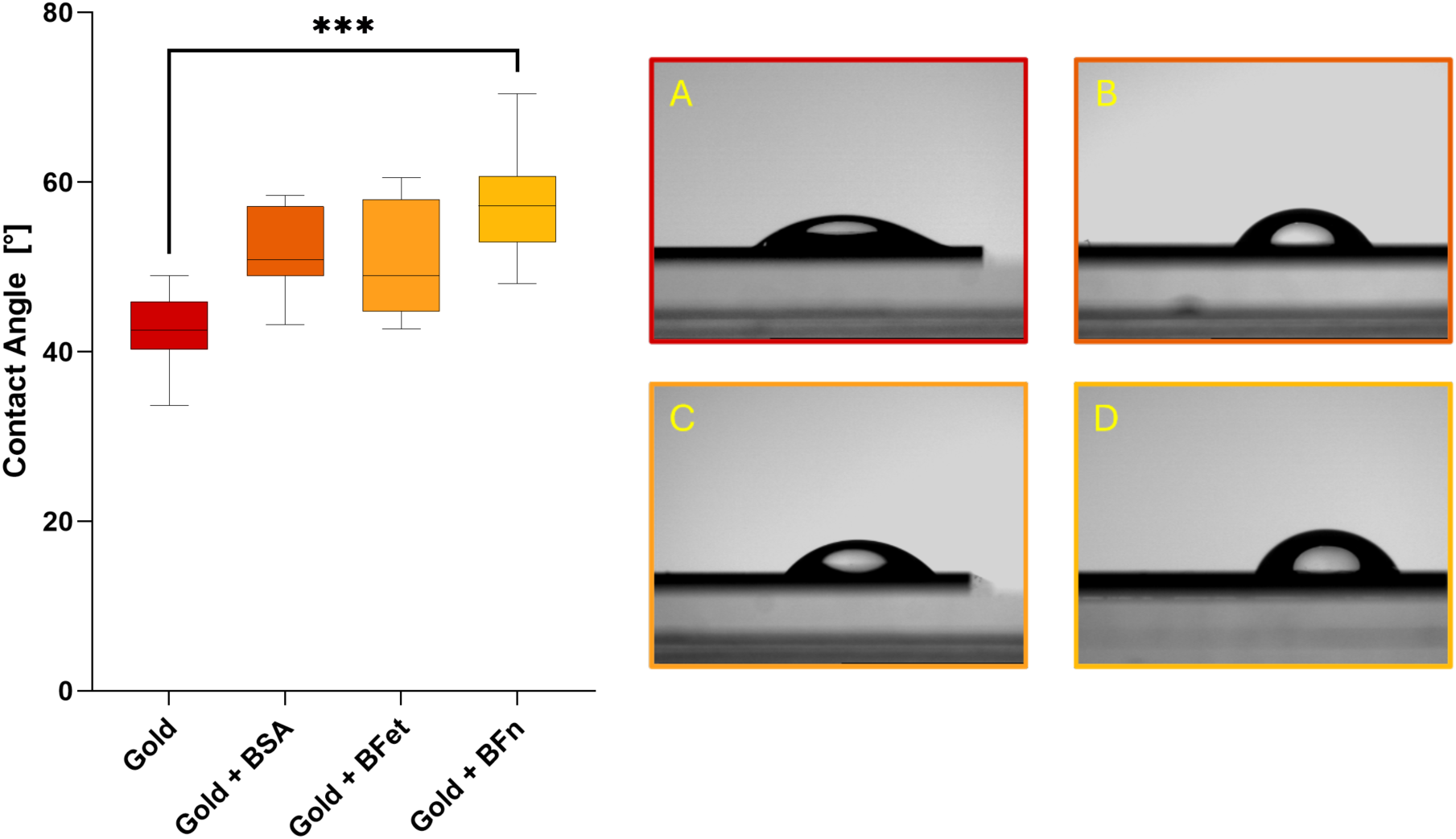
Water contact angle measurements obtained through the sessile drop method. Results are presented as average ± SD, N ≥ 12. On the right side, representative images of contact angle droplets for Gold (A), Gold + BSA (B), Gold + BFet (C) and Gold + BFn (D) are presented.

Gold surfaces in typical air exposed environments generally exhibit moderate water contact angle measurements, in the range of 60-65° ^50,51^. The lower contact angles detected for the cleansed gold surfaces attests to an effective cleaning of the surface through the treatment in TL1 solution, and likely elimination of most organic contaminants present on the surface, in line with previous studies ^50,52,53^. While an increase in contact angle was detected upon the adsorption of proteins on the model surfaces, no significant difference was found among the different protein modified samples, with the exception of BFn, whose presence on the surface resulted in a CA increase around 10° higher than other proteins.

### Correlative Protein Quantification

When an orthopedic implant is inserted into the body, the proteins present in the physiological fluids readily adsorb to the surface, and biomolecules re-arrange their three-dimensional structure at the solid-liquid interface by forming secondary bonds and potentially exposing certain motifs along their chain. The surface is quickly saturated by the binding of lighter, more abundant proteins, which form at least a protein monolayer at the materials interface. The composition and characteristics of such protein layer(s), however, evolves over time through a process called the Vroman effect, where higher concentration, lower-affinity molecules desorb and are substituted by other lower concentration but higher-affinity proteins^54–56^.

The BSA protein used for adsorption comparison, presents a similar molecular weight to BFet (around 66 kDa for albumin^37^ and around 60 kDa for fetuin-A, but varying slightly depending on the degree of glycosylation^23^). They also have a similar structure in solution, albumin has a globular structure with reported hydrodynamic radius of 3.51 nm^38^, while fetuin-A presents a globular structure with hydrodynamic radius of around 4.2 nm, but an increase in size of around 10% upon interaction with Ca/P ions ^57^.

Four different techniques were employed to address and correlate the protein interactions on gold surfaces. These provided adsorption quantification, allowed comparison of BFet to BSA, binding kinetics, changes in adsorbed layer characteristics, and remaining protein levels after treatment with SDS surfactant to elute loosely bound protein.

Representative curves for frequency (in orange) and dissipation (in green) obtained with QCM-D for both proteins are shown in Figure 2A, and provide insight into the different interactions with water molecules between the two proteins. Over the 3 h adsorption, saturation of the surface coverage is reached within the first 20 min from injection. The buffer-assisted rinse (PBS) under equal flow conditions, performed to remove any loosely-bound protein from the surface, negligibly decreased the quantity of protein present on the surface, which retained more than 97% of the adsorbed mass for both proteins, indicating a stable adsorption of both proteins to the substrate. The mass adsorbed measured through QCM-D at plateau conditions (Figure 2B, solid lines) is however notably different between the two proteins (882 ng/cm^2^ for BFet and 540 ng/cm^2^ for BSA, respectively). Association constants, as measured through QCM-D, were 18553±2755 min^-1^ M^-1^ for BSA and 56397±1620 min^-1^ M^-1^ for BFet.

**Figure 2.**
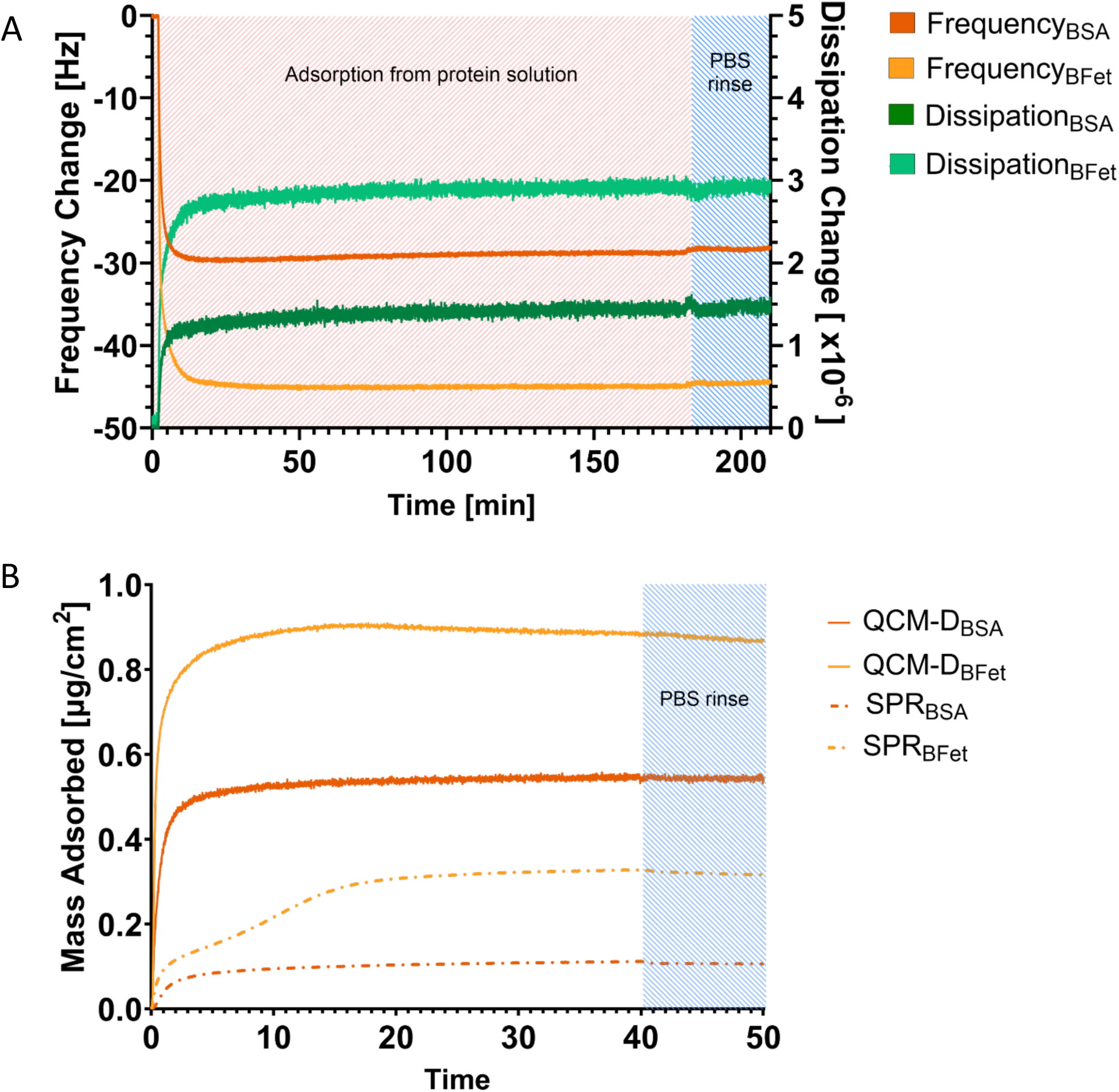
(A) Comparison of frequency (orange) and dissipation (green) changes observed in quartz crystal microbalance with dissipation (QCM-D) upon adsorption of BSA and BFet. The higher measured dissipation for BFet indicates a less compact and more hydrated adsorbed protein layer with respect to BSA. (B) Comparison of measured mass adsorbed to the surface at saturation detected through QCM-D and surface plasmon resonance (SPR). Solid lines represent results obtained through QCM-D, 5^th^ overtone, which measures the hydrated weight of the adsorbed protein; dotted lines represent the net weight of adsorbed protein as measured through SPR. BSA is presented in dark orange, BFet in light orange.

The adsorbed surface density at plateau conditions measured through SPR after 50 min from injection is presented for both proteins in Figure 2B (dotted lines), with BFet displaying a surface coverage of 270±52 ng/cm^2^ and BSA 109±4 ng/cm^2^. Note, the mass coverages measured by SPR correspond to the pure protein coverage, whereas the mass coverage measured by QCM-D also includes any associated water with the protein. The amount of protein adsorbed was confirmed with complementary dry SPR measurements after 180 min submersion of the samples in 0.2 mg/mL protein solution, PBS washing and nitrogen drying, showing similar mass coverages. The resulting layer thickness and calculated mass adsorption are presented in Figure 1S (I and II).

Protein quantification obtained through radiolabelling (Figure 3) confirms similar adsorption masses for BFet with respect to BSA. Longer elution time in the presence of the surfactant, combined with shaking conditions, led to a significant increase in the amount of protein eluted from the surface. However, while both proteins retained around 75% of the originally adsorbed mass, the surfactant caused a higher amount of BFet to come off the surface with respect to BSA. Sodium dodecyl-sulfate can disrupt non-covalent bonds, such as hydrogen bonds, or hydrophobic interactions, but by itself cannot break disulfide bonds. The higher elution of BFet from the gold surface could be related to the lower amount of disulfide bonds throughout the molecule (6 internal disulfide bonds for fetuin opposed to the 14 for albumin) available for interaction with the gold surface, or a possible difference in binding mechanism to the surface with preference toward hydrogen bonds or hydrophobic interactions.

**Figure 3.**
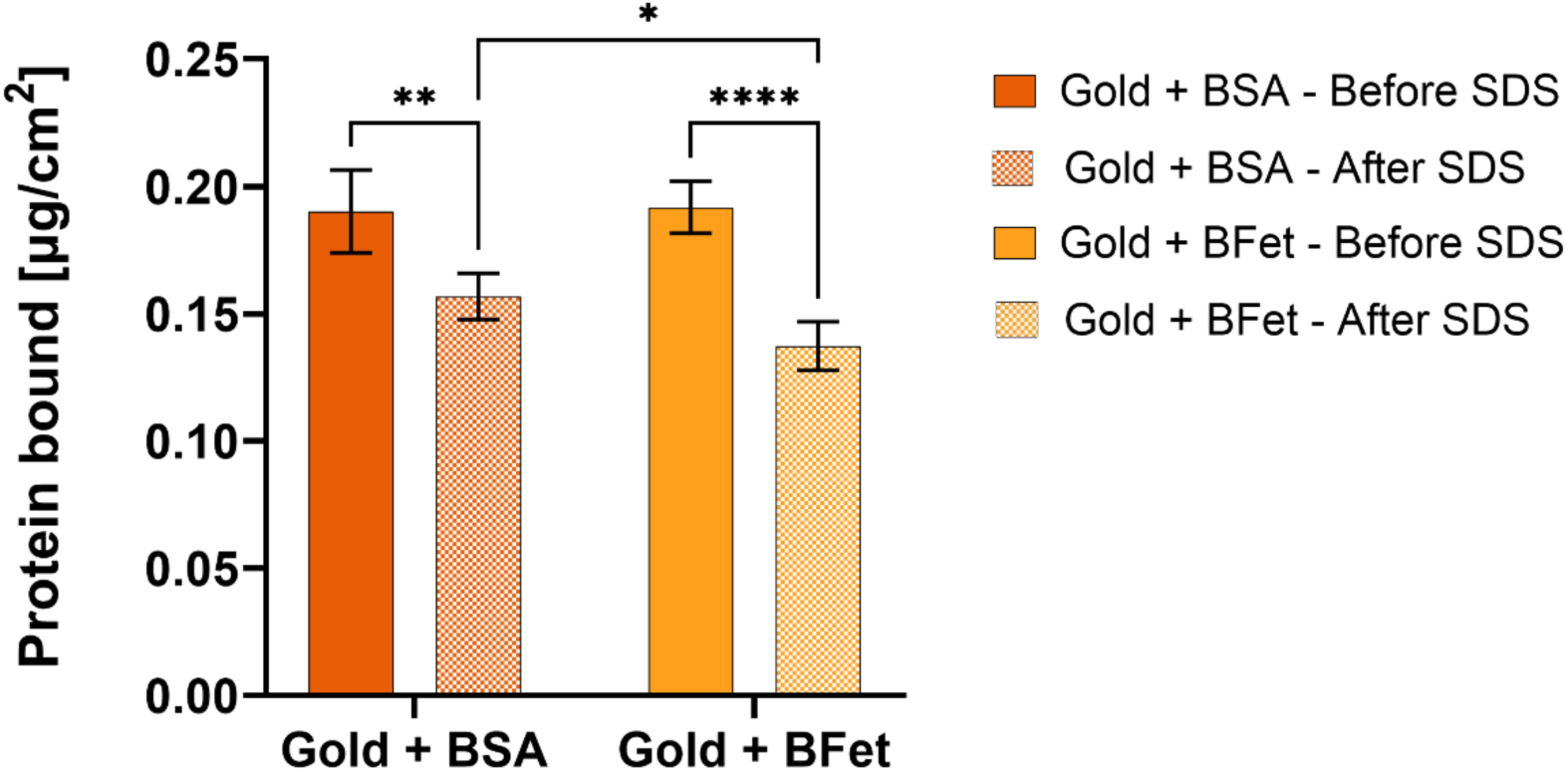
Protein mass adsorbed as measured by protein radiolabelling with ^125^I. Values are normalized over the surface area after 3 h of adsorption under static conditions (before SDS) and following overnight elution in 2% SDS (after SDS, textured columns). Results are presented as average ± SD, N=4. BFet binds in similar amounts as BSA to gold surfaces, with a total adsorbed mass of around 0.2 µg/cm^2^ after 3 h in 0.2 mg/mL protein solution.

It should be noted that the protein quantification results obtained from SPR and QCM-D for BSA, and further confirmed by the radiolabelling results, are below the commonly accepted calculations for the protein mass for monolayer formation based on molecular weight and hydrodynamic radius of the protein. The values obtained suggest instead a notable re-arrangement of the BSA 3D structure upon binding to the gold, moving from a globular conformation in liquid to a more spread-out arrangement at the interface. Such mass adsorption is in line with previous observations using analogous conditions: Tworek et al^58^ investigated the adsorption effectiveness and structural rearrangement of BSA upon adsorption on gold surfaces. At physiological pH and ionic strength similar to that of PBS, BSA adsorbed on the gold surface with a mass coverage of ∼ 100 ng/cm^2^, in line with the present study, adopting a mixture of elliptical and triangular orientations for the formation of the monolayer. The group also observed a change in distribution of secondary structures of the protein upon adsorption at the interface, mainly composed of α-helixes and β-turns. Such values for BSA are aligned with the results obtained in the present investigation for mass adsorption, where BSA coverage is around 109 ng/cm^2^. In comparison, however, BFet adsorption amounted to around 270 ng/cm^2^, a value more in line with the expected density of a fetuin monolayer. Such discrepancy between BSA and BFet, with all other parameters constant, points toward a difference in structural re-arrangement of the two proteins upon interaction with the substrate: a more “spread-out” conformation for BSA with respect to a higher density packing for BFet, with less 3D re-arrangement of the latter upon interaction with gold. Such difference between BSA and BFet was also observed in the complementary SPR dry measurements: higher thickness and adsorbed mass were measured for BFet with respect to BSA (Figure S2, I and II), pointing toward an overall higher fetuin-A adsorption from equally concentrated solutions, with limited re-arrangement upon interaction with the surface and retainment of a more compact, less spread, 3D structure.

The mass adsorbed at plateau conditions measured through QCM-D was also higher for BFet, suggesting a higher affinity for water binding and entrapment in the BFet adsorbed layer with respect to the BSA layer. Such results were also supported by the difference in dissipation curves between the two proteins, which could indicate a different re-arrangement of the adsorbed layer between the two. The differences in amino acid sequences between the two proteins, and thus the difference in number or strength of intermolecular bonds, as well as in the number and strength of bonds between the molecule and the material, may explain the difference in 3D re-arrangement upon binding.

The comparative mass adsorptions determined from the three correlative techniques of SPR, QCM-D and radiolabeling present similar trends to what has been previously observed^59^, with SPR and radiolabeling reporting similar values for adsorption, accounting for the dry weight of the mass adsorbed, with QCM-D measuring “wet” weight, also including the component due to water entrapped in the layer, thus overestimating the actual amount of protein bound to the surface. A comparison of the results obtained using the three methods is presented in Figure 1S, III.

Quantitative confirmation of homogeneous surface coverage is provided by fluorescence microscopy results of fluorescently-tagged protein distribution on gold surfaces. Both gold+BSA (Figure 4A) and gold+BFet (Figure 4B) present a homogenous surface coverage, quantified by the <CV> of the images for each sample, with <CV_BSA_> = (13.6 ± 0.2)% and <CV_BFet_> = (13.4 ± 0.2)%. From the images, the fluorescence intensity of BFet, normalized over the inherent fluorescence of the gold surface and accounting for each protein’s DOL, is approximately 2.5 times that of BSA on gold surfaces, providing further confirmation of the higher amount of BFet mass adsorbed to the gold surface compared to BSA. The image processing procedure for fluorescence results quantification is presented in Figure 2S.

**Figure 4.**
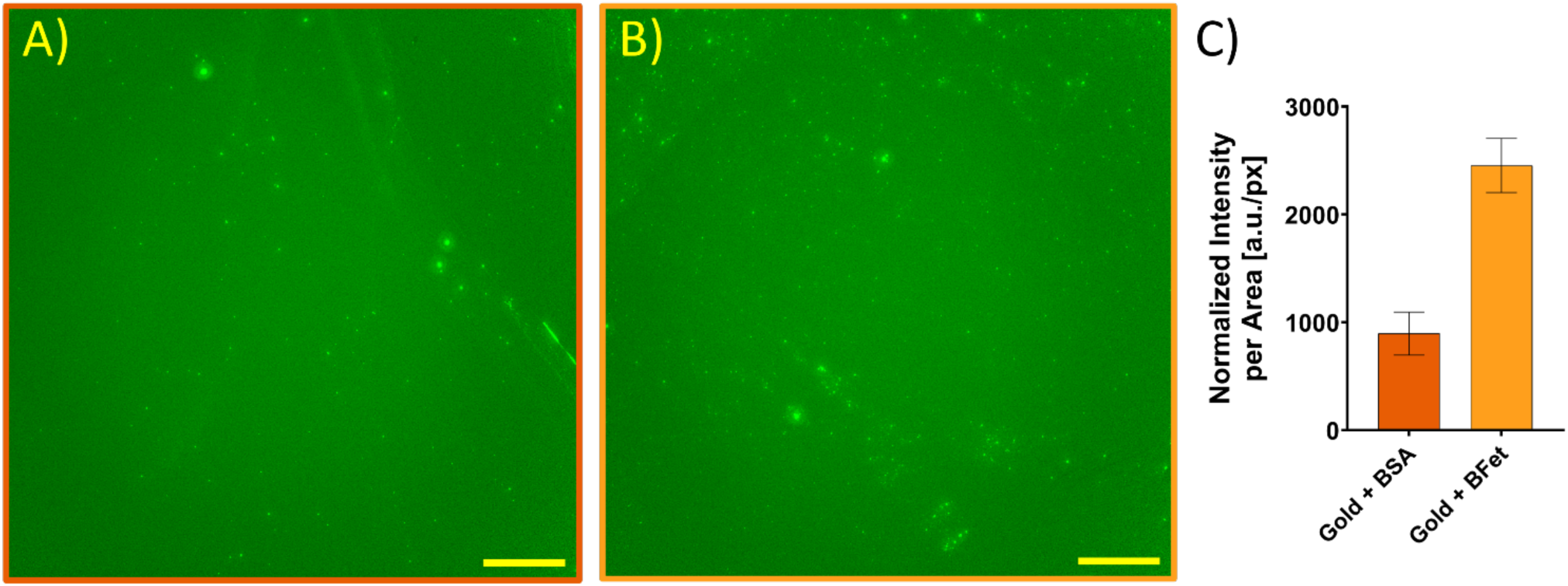
Representative fluorescence microscopy images of gold substrates functionalized with either (A) fluorescent BSA or (B) fluorescent BFet. Scale bars are 20 µm in length. Fluorescent proteins were very evenly distributed across gold substrate surfaces, with only some aggregates present. (C) The normalized, average fluorescence intensity measured per pixel for BSA and BFet on gold.

Protein bioactivity depends upon the availability of the protein’s bioactive sites towards their substrates. When proteins aggregate, these bioactive sites can be hidden from the substrates, diminishing their bioactivity ^60,61^. As protein surface adsorption can influence aggregation ^60^, quantitative analysis of BSA and BFet aggregates on gold substrates provides information on the potential bioactivity of these surfaces. Using the Max Entropy mask described above, the percent surface area (% SA) occupied by aggregates for each protein was found to be very minimal, with % SA_BSA_ = 0.2 ± 0.3% and % SA_BFet_ = 0.3 ± 0.3%, indicating that over 99% of the gold surfaces are occupied by non-aggregated proteins in both cases. To further quantify the protein surface adsorption across the gold substrates and ensure that both proteins were similarly distributed, we calculated the CV of the images, excluding the aggregates. In the fluorescent images with the aggregates excluded, the CV values averaged across all images for each protein were found to be ∼13.5%. The high similarities between these CV values suggest similar adsorption mechanisms to gold and distributions across the surface of the substrate. The low CV values (close to 0%) also show a uniform protein distribution across the gold substrate surfaces, indicating that the surface area excluding the aggregates is nearly uniform with protein that likely retains its bioactivity.

### Cellular Response Analysis

To address the adhesive capabilities of fetuin-A, its role in cellular attachment and proliferation of osteoblast-like cells was assessed through fluorescence imaging of fixed and stained cells, and interpreted based on changes in cellular metabolism detected through a fluorescence assay. To prevent possible substitution of the pre-adsorbed proteins for cell-adhesive proteins present in fetal bovine serum, due to the Vroman Effect^62^, cells were first seeded in non-supplemented media^63^. Preliminary studies confirmed that there was no significant effect on cellular metabolism from the use of non-supplemented media for the first 24 h of culture, followed by 15% substitution with supplemented media for the remaining time, as per the suggestions of the cell provider to sustain cell growth and proliferation (Figure 3S). In this 24 h time-frame, cells are then only able to attach to the surface through the pre-adsorbed proteins or other mechanisms, and not through the cell adhesive moieties present in some proteins that constitute FBS.

To assess cellular attachment, cells were fixed and stained for nuclei and actin filaments: nuclei were counted to detect differences in the number of cells attached based on pre-adsorbed protein after 24 h, and image analysis was used to identify discrepancies after 24 h from seeding in relevant cell parameters such as cell eccentricity, cell area, and cell major and minor axis, descriptive of the specific phenotype. Cellular proliferation was assessed by evaluating the nuclei count over 7 days, as well as the total surface coverage of cells.

No statistically significant difference was found in number of cell nuclei on the surfaces with adsorbed proteins after 24 h from seeding (Figure 5). Nonetheless, the average number of cells on BFn and BFet both exceed the control surface value, with BSA being the only pre-adsorbed protein presenting a lower nuclei count compared to the bare gold surface. Fibronectin’s amino acid sequence presents the cell-adhesive RGD tripeptide, known to promote cell attachment and proliferation through integrin binding. The results obtained on pre-adsorbed BFet surfaces, comparable to those of the positive BFn protein control, point thus toward cell adhesive capabilities of fetuin-A, in agreement with previous investigations^64,65^. On day 3, cells on gold+BSA behaved comparably, on average, to cells on gold. In contrast, cells attached to gold+BFet demonstrate a higher cell nuclei count than the surface control and negative BSA protein control but are lower than the number of cells on gold+BFn. Such findings suggest that BFet, as opposed to BFn, does not play an active role in the promotion of cellular proliferation. All surfaces at confluency present a similar cell nuclei count, excluding any effect from the protein. Representative fluorescence micrographs used for cell counting over 7 days are provided in Figure 4S.

**Figure 5.**
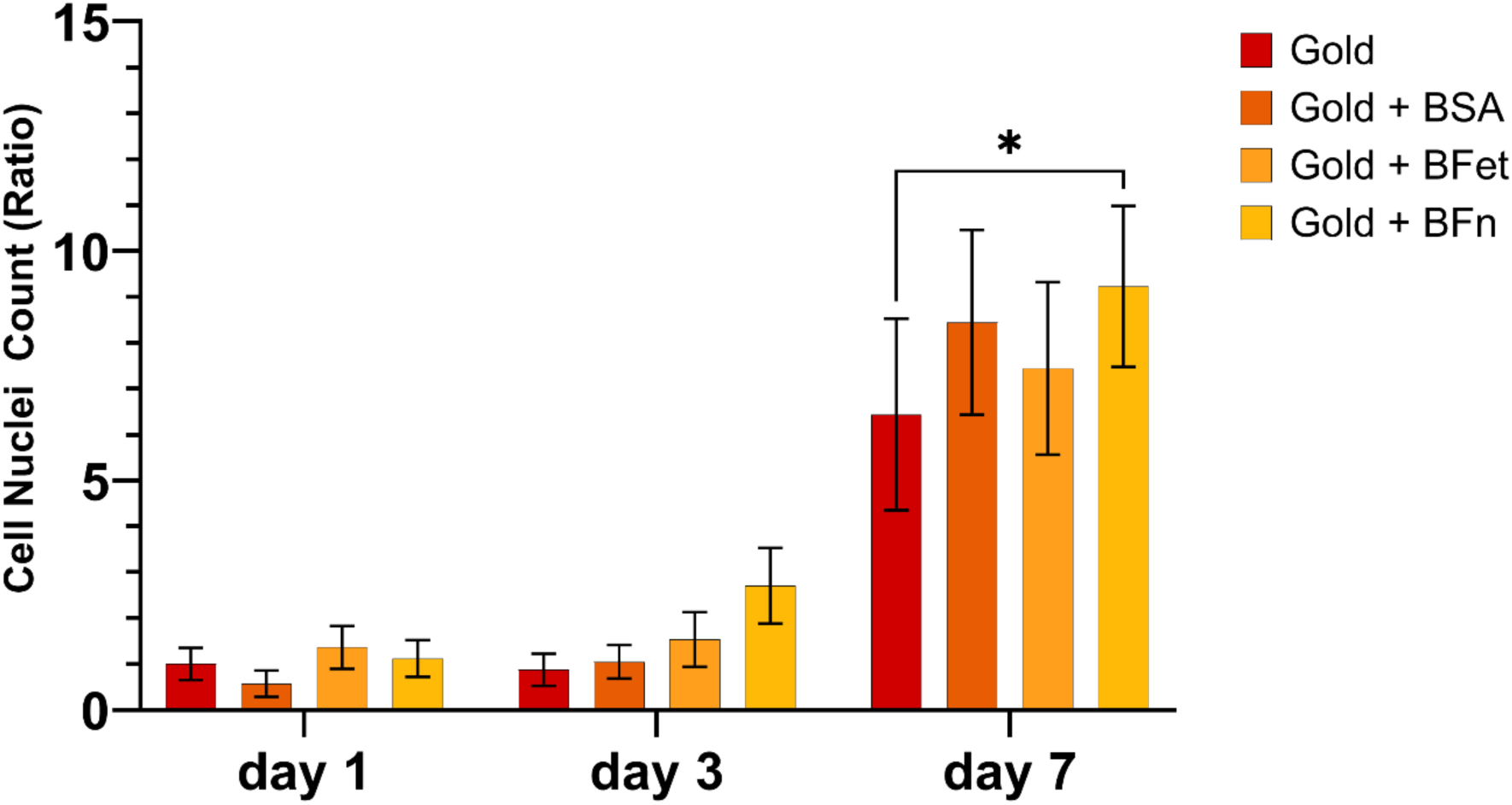
Ratio of cell nuclei number on either the bare gold or pre-adsorbed protein surfaces at different time points (day 1, 3, 7). Values are normalized over the average value of the number of nuclei on gold at day 1. Results are presented as average±SD. No statistical difference was found between samples on the first day, but on average BFet performs comparably to the positive control BFn, suggesting an active role in cellular attachment. BFet does not present an active role in the sustainment of cellular proliferation over 7 days.

The differences observed in number of cells attached over 24 h were further assessed through the image analysis of single cells grown on sample surfaces to evaluate morphological characteristics. While BFet seemed to enhance cellular adhesion based on cell nuclei count, similarly to BFn, the results from day 1 of the morphological analysis present differences between BFet and BFn. The average surface area for single cells on all substrates is close to 1000 µm^2^, as expected from previous findings^46,66^. On gold and gold+BFn, however, the size of individual cells (cell area) was higher with respect to gold+BSA and gold+BFet (Figure 6A), while for the eccentricity (Figure 6B), the average value for cells on bare gold is noticeably lower than the surfaces presenting adsorbed proteins. The major axis results (Figure 6C) do not present relevant differences among the adsorbed surfaces, apart from BFn, which has a significantly higher major axis when compared to the other surfaces. These findings are further confirmed by the comparison of the cell minor axis among the different conditions (Figure 6D). Cells attached on pre-adsorbed BFet gold surfaces present, in conclusion, similar eccentricity, but smaller surface area and major axis length than those on BFn. These differences indicate cells are well-adhered to the surface and present morphological epithelial-like characteristics expected from the cell line, but are less spread out than cells on BFn. Even if BFet promotes cell attachment, the morphological characteristics of the cells grown on it are more similar to those of cells on gold+BSA surfaces. Cells that adhered to gold only, in the absence of serum proteins, present a surface area and min axis equal to those on gold+BFn, but lower eccentricity and max axis. Given the characteristics of the cell-line in use, a more circular morphology is indicative of a condition that is not beneficial for cell attachment and proliferation.

**Figure 6.**
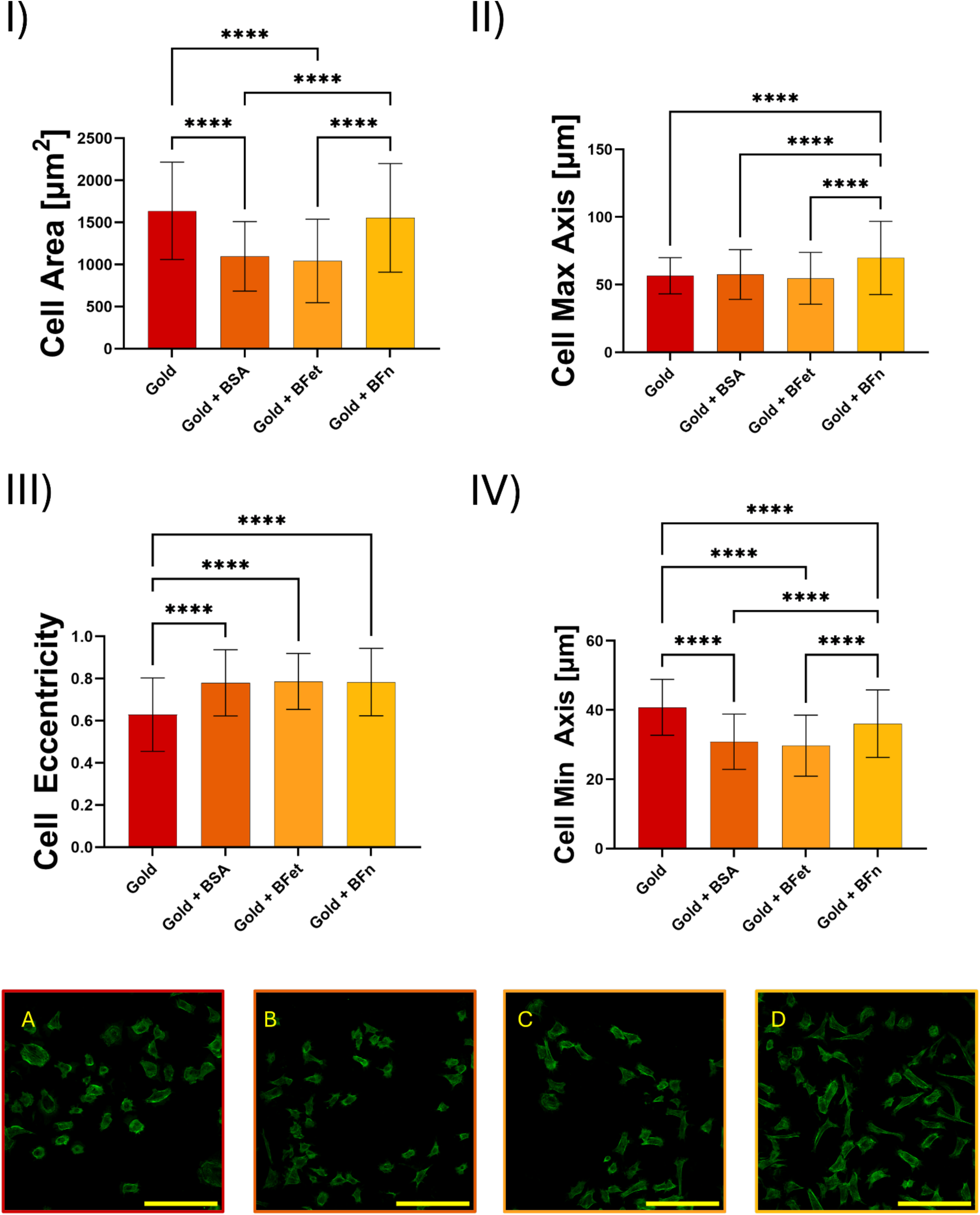
Differences in relevant cell morphological characteristics on either the bare gold or pre-adsorbed protein surfaces at day 1 as measured through surface image analysis of single-standing cells: I) cell area, in µm^2^, II) cell eccentricity, III) size of major axis, IV) size of minor axis. Results are presented as average±SD. Representative images displaying morphological differences in cells attached on A) Gold, B) Gold+BSA, C) Gold+BFet and D) Gold+BFn are presented in the fluorescence micrographs. Scale bar is 200µm. Saos2 cells attached on gold+BFet surfaces present morphological characteristics similar to those on gold+BSA, with similar eccentricity but lower cell area and max cell axis with respect to the positive control gold+BFn.

The results for cell proliferation based on nuclei count over different time points were corroborated by the percentage surface coverage results (Figure 7). For day 1, no statistically significant difference was found in terms of the percentage of substrate surface covered by cells. Nonetheless, both BFet and BFn present higher surface coverage with respect to the bare gold surface. On day 3, a statistically significant increase in surface area covered by cells can be observed for the pre-adsorbed BFn surfaces, whose cell surface coverage almost doubles, on average, the other surfaces’ results. All other adsorbed conditions demonstrate similar cell behaviour compared to the gold control, with a surface coverage around 30%. After 7 days, all substrates presented comparable surface coverage by cells with a confluent layer.

**Figure 7.**
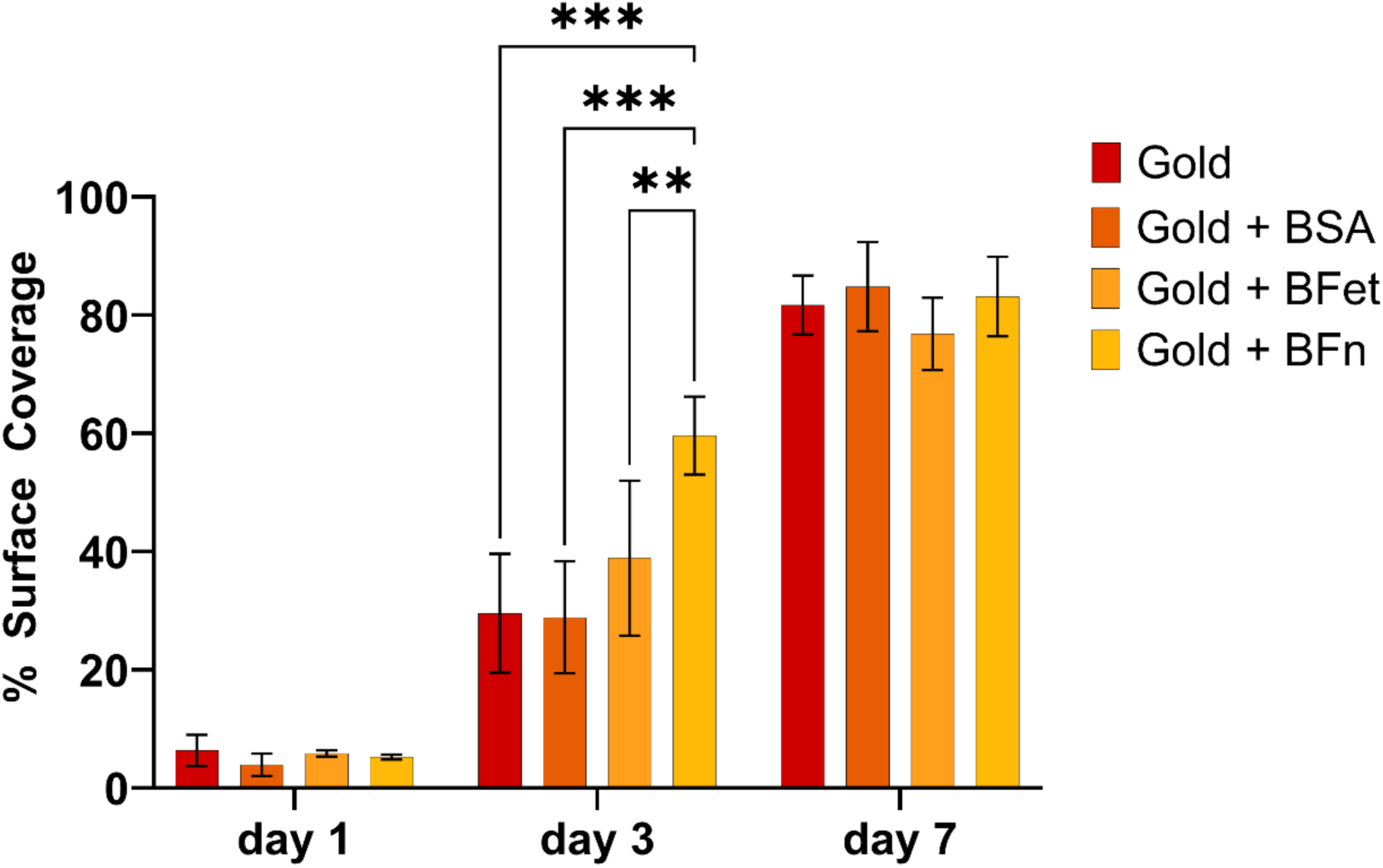
Percentage area of the sample covered by cell bodies on either the bare gold or pre-adsorbed protein surfaces, at three different time points (day 1, 3, 7). Results are presented as average±SD. BFet seems to increase % cell surface on day one, likely because of higher average number of cells attached, but displays no role in the sustainment of cellular proliferation over 7 days.

The results from total surface coverage, thus, support the observations from the cell nuclei count. On day 1, gold+BFet performed analogously to gold+BFn, supporting the hypothesis that fetuin-A plays an active role in enhancing osteoblast-like cell attachment. On day 3, cell growth on gold+BFn outperformed that on all other surfaces, demonstrating BFn’s ability to enhance cellular proliferation and greater cell spreading. While lower than that on gold+BFn, cell growth on gold+BFet substrates exceeded, on average, that on gold and gold+BSA. Yet, no active enhancement of cellular proliferation could be detected.

To complement the observations on cell attachment and proliferation, cell metabolism was assessed using an AlamarBlue assay at three different time points (Figure 8) and correlated with the results from immunostaining of the cells grown on substrates with DAPI and Phallodin-Alexa488 at the same time points. The AlamarBlue assay provides information on the viability and proliferation of cells attached to a substrate through the transformation of resazurin to fluorescent resorufin produced by metabolically active cells. Upon attachment, no statistically significant difference was found among the different conditions. However, on average, cells grown on gold+BFet behave similarly to those on bare gold and gold+BFn, while those grown on pre-adsorbed BSA present lower metabolism.

**Figure 8.**
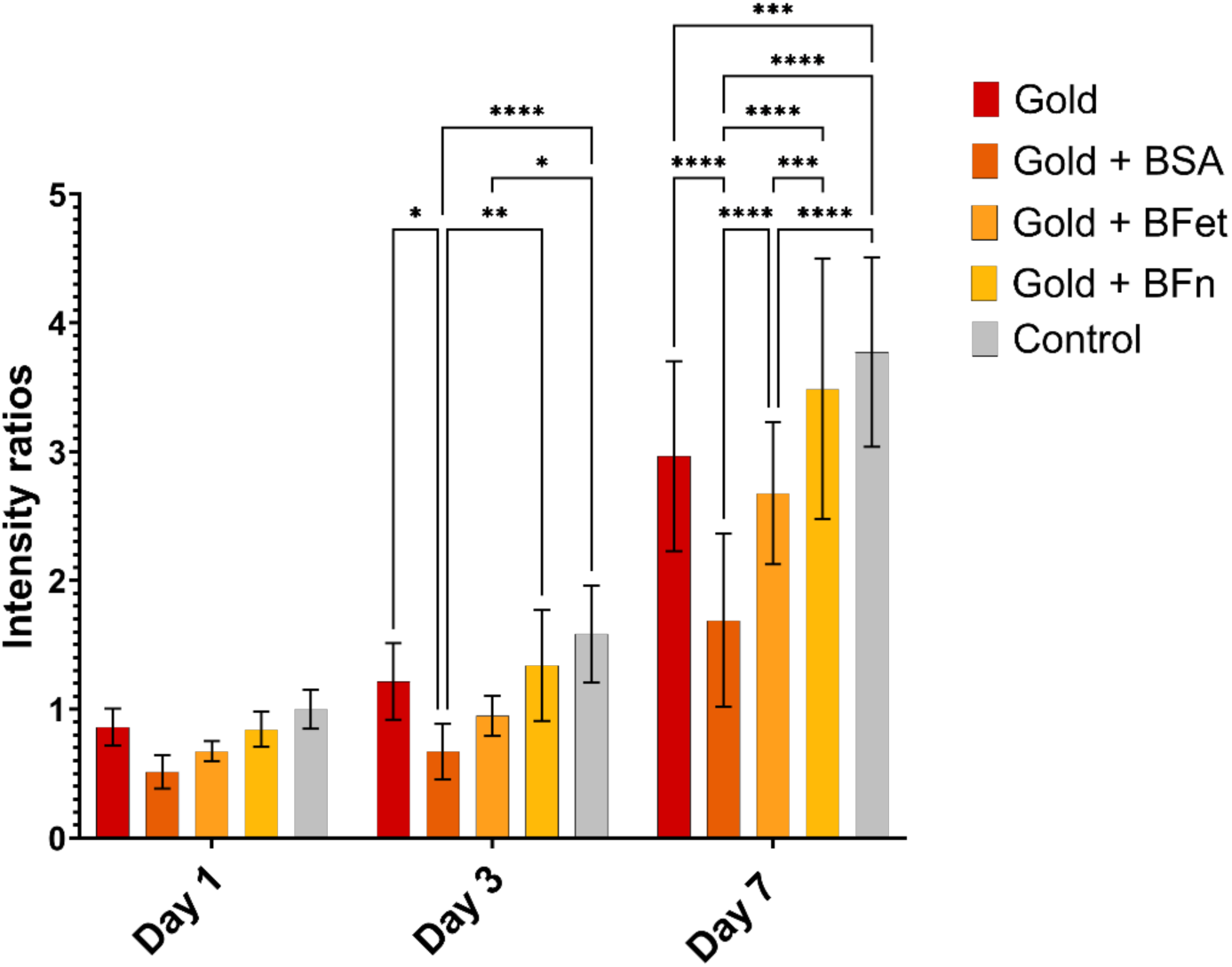
Fluorescence intensity values representing overall cell metabolism for cells grown on either bare gold or pre-adsorbed protein surfaces at different time points (day 1, 3, 7). Values are normalized over the average intensity value of the polystyrene plate control at day 1. Results are presented as average±SD. Cells attached on gold+BFet, while presenting similar metabolism on day 1 to the cells on gold and gold+BFn, display a decrease in cellular metabolism over 7 days with respect to the positive control (BFn).

On day 3, however, cells attached to gold+BSA substrates presented a significant decrease in fluorescence intensity compared to that of cells on gold, gold+BFn, and the tissue culture polystyrene (TCPS) plate (control). In contrast, no statistically significant difference in cell metabolism was detected for cells grown on BFet compared to BFn and BSA, whereas cells cultured on gold+BFet presented lower metabolism compared to the TCPS control. The absence of significant differences between the metabolism of cells grown on gold+BFn and the control confirms that a decrease in metabolism is not due to a change in the substrate chemistry but rather depends on the protein properties themselves. Such findings agree with observations by Oschatz et al on a different osteoblast-like cell line^36^, where a similar change in metabolism for cells grown on fetuin-A coated surfaces was detected over 48 h.

Day 7 results provide further confirmation of the decrease in metabolism for Saos2 cells cultured on gold+BFet, with a statistically significant decrease in fluorescence intensity compared to the bare gold, gold+BFn, and TCPS substrates. However, the increase in metabolism for cells grown on substrates with BFet is statistically significant compared to the negative protein control, gold+BSA. In conclusion, this reduction in metabolism for cells on substrates pre-adsorbed with BFet compared to the positive control may be linked to the absence of enhanced cellular proliferation on these same substrates, as reduced metabolism will affect proliferation.

It should be noted that, upon exchanging non-supplemented media with supplemented media after 24 h, the composition of the adsorbed protein layer on the surface may have changed with respect to day 1. As such, the cell proliferation results for days 3 and 7 could be partially due to other proteins within the supplemented media adsorbing to the surface, replacing the pre-adsorbed proteins, or the interaction of the pre-adsorbed protein with other biomolecules or ions present in serum. This applies particularly to bare gold surfaces, where the variation in the substrate chemistry following protein adsorption could explain the higher cell metabolism measured over 7 days for gold surfaces and good standing of the cells, in spite of the rounder morphology presented after the first 24 h. However, over the first 24 h, cell-binding to the surface was possible only through specific interactions with the proteins pre-adsorbed on the surface, or through non-specific interactions, as is most likely in the case of bare gold. In addition, while notable differences exist in the measured metabolic activity over 7 days among the different samples, no relevant differences were observed between the samples in terms of the number of cells present on the surface. This supports the idea that pre-adsorbed fetuin-A likely plays a role in cellular metabolic pathways.

Plasma fetuin-A has been found to act as an antagonist of transforming growth factor-β (TGF-β) and other bone morphogenic proteins^67,68^, both of which play a role in bone formation and metabolism. Fetuin-A is also reportedly involved in tumor growth and metastasis, with contrasting reports on its pro-or anti-tumor development potential based on the organs involved^69–71^. In the case of bone tumors, fetuin-A was reported to attract tumor cells, a characteristic which could explain the increased cellular attachment observed both in the present investigation and in the study by Oschatz et al^36^. In contrast, plasma fetuin-A was previously reported to increase cell growth and proliferation^72,73^, specifically for carcinogenic cell lines^69,74–76^, whereas in both investigations addressing fetuin-A pre-adsorption on a biomaterial interface, a decrease in cellular metabolism was observed. The present investigation also detected, in contrast to previous reports ^36,65,77^, a decrease in proliferation over 72 h for cells in contact with fetuin-A when compared to a protein with cell-adhesive properties. It is worth noting that in previous work by Vyner and Amsden^65^, the improvement of cellular proliferation related to pre-adsorbed fetuin-A was hypothesized to be connected to a synergistic effect between fetuin-A and other proteins and growth factors present in plasma. This work supports the hypothesis that fetuin-A, alone, may not be able to sustain cell proliferation. In physiological systems, over a hundred fetuin-A molecules bound to calcium phosphate (CaP) in the bloodstream form 100-nm large aggregate CPPs, with the CaP’s form being either amorphous or hydroxyapatite. As the CaP form itself can impact cellular processes, biomaterial surface modifications involving fetuin-A may be most effective at influencing bone cell behavior when they are adsorbed as pre-formed CPPs. Further investigations are needed to fully clarify the effects of fetuin-A pre-adsorption at the biomaterial interface on subsequent bone cell proliferation and mineralization, and to address any differences between physiological and pathological conditions, as well as the specificities related to cell lines and substrate material properties.

## Conclusions

The results of this study demonstrated that fetuin-A adsorbed on model gold surfaces with similar kinetics to those of albumin, but in higher amounts and with a more hydrated layer, as highlighted by the QCM-D results. The SPR and fluorescence microscopy observations suggested the formation of a protein monolayer on the surface, with limited 3D re-arrangement of fetuin upon adsorption when compared to a more spread-out structure adopted by albumin, as previously observed in literature. In addition, when pre-adsorbed on the surface, fetuin-A slightly increased the binding of Saos2 cells, in line with previous observations of other cell lines. However, the morphology of the osteosarcoma cells presented lower surface area and elongation when compared to surfaces coated with fibronectin. At longer time points, the pre-adsorbed protein did not present any beneficial role in cellular proliferation, likely related to the decrease in cellular metabolism detected for pre-adsorbed fetuin-A surfaces. In conclusion, fetuin-A has the potential to be used as a surface modifier for improved cellular attachment to a surface, but by itself does not, alone, improve the proliferation of osteoblast-like cells in the early stages of the osteointegration process, likely due to a decrease in cellular metabolism. Further investigations are needed to address the specificity of such findings related to the substrate characteristics, both chemical and topographical, and the particular cell lines.

## Acknowledgements

Confocal microscopy and *in vitro* studies were carried out at the Centre for Advanced Light Microscopy and the Biointerfaces Institute at McMaster University.

Dr. João Bronze de Firmino, Dr. Bryan Lee, Dr. Joseph Deering and Dr. John Andersson are acknowledged for scientific discussions and support.

K.G. and K.N.S. acknowledge funding support from the NSERC Discovery Grant Program (grants RGPIN-2020-05722, RGPIN-2019-06433, and RGPIN-2022-05258, respectively), Mitacs Accelerate Program and Mitacs Globalink Research Abroad Program (Application Ref. IT27778 and IT39044, respectively), the New Frontiers Research Fund and Mitacs, and K.G. from the Canada Research Chairs Program (Tier II Chair in Microscopy of Biomaterials and Biointerfaces). A.M. acknowledges the funding and support of the Foundation Blanceflor Boncompagni Ludovisi, née Bildt, and Mitacs Accelerate Program. BioRender was used for the graphical abstract (license FP28UIAY61).

## Author Contributions

**Alessandra Merlo**: conceptualization, investigation, methodology, data analysis, image analysis, visualization, manuscript – original draft. **Jesper Medin**: investigation, data analysis. **Shane Scott**: investigation, data analysis. **Andreas Dahlin**: conceptualization, supervision, funding. **Kathryn Grandfield**: conceptualization, supervision, funding. **Kyla Sask**: conceptualization, supervision, funding. All authors reviewed and edited the present manuscript.

