## Supporting Information for "The Role of Fetuin-A on the Attachment and Proliferation of Osteoblast-like Cells on Model Gold Surfaces"

**Section 1S. Additional Methodology Description**

**1S.1 Addition to Section 2.4 of Materials and Methods: Quartz-Crystal Microbalance with Dissipation**

The association constants for both proteins were obtained using the Langmuir adsorption model (Equation S1):

Equation S1

$$\frac{d\Gamma}{dt}=k_{on}C\left( t \right)\left[ \Gamma_{max}-\Gamma\left( t \right) \right]-k_{off}\Gamma(t)$$

Where Γ indicates the fraction of occupied adsorption sites, k*_on_* indicates the association constant, C indicates the concentration of the adsorbate, and k*_off_* the dissociation constant.

Assuming C(t) is constant at C_0_, Equation S1 can be rewritten as Equation S2:

Equation S2

$$\frac{d\Gamma}{dt}=k_{on}C_{0}\left[ \Gamma_{max}-\Gamma\left( t \right) \right]-k_{off}\Gamma(t)$$

Assuming that Γ(t) $\approx$0 for small t values, Equation S2 through can be rewritten as Equation S3:

$$\left. \frac{d\Gamma}{dt} \right|_{0}=k_{on}C_{0}\Gamma_{max}$$

Equation S3

the association constant can be calculated considering the value of $\left. \frac{d\Gamma}{dt} \right|_{0}$ equal to the initial slope, C_0_ as the initial protein concentration, and Γ_max_ as the maximum surface coverage as measured through the wet 50 min injection SPR measurements.

**1S.2 Addition to Section 2.7 of Materials and Methods: Fluorescently-tagged Protein Surface Quantification**

1S.1 Fluorescent Protein Quantification

To determine protein concentrations and their degrees of labelling (DOL), the number of dyes per protein molecule, were determined by measuring their absorbance on a Cary 60 spectrophotometer (Agilent, Santa Clara, CA, USA). Protein concentration was quantified using the extinction coefficients for BSA and BFet provided by Sigma Aldrich and calculated using Beer’s law. The DOL for each protein was calculated using

$$DOL = \frac{Abs\left( 495 nm \right)}{\varepsilon_{Alexa488}}\cdot\frac{\varepsilon_{protein}}{Abs\left( 280 nm \right)}$$

where Abs(λ) is the absorbance at wavelength λ, and ε_x_ is the extinction coefficient for x = Alexa488, or protein. BSA was found to have DOL = 0.802, while BFet had DOL = 0.491.

Equation S4

1S.2 Fluorescence Microscopy Image Quantification

To calculate the normalized average fluorescence intensity, the average pixel intensity of each image was calculated using Fiji. From the average pixel intensities of each image of the control, BSA, and BFet samples, the mean pixel intensity for each data point was calculated, with the standard deviation between the average pixel intensities for a given sample calculated. From these values, the normalized average fluorescence intensity <I_protein, norm_> for each protein-functionalized gold sample was calculated by the formula

Equation S5

$$\left\langle I_{protein, norm} \right\rangle=\frac{\left\langle I_{protein} \right\rangle-\left\langle I_{Au Control} \right\rangle}{DOL}$$

In Equation S2, <I_protein_> is the average pixel intensity of all the images for each protein, <I_Au Control_> is the average pixel intensity of all the images of the control, and DOL is the degree of labelling of the protein. As the overall measured fluorescence intensity per pixel is a linear combination of the contribution from the light reflected from the gold surface, regular background light, and emission by fluorescently-labelled proteins, the light intensity contribution from the proteins can be calculated. This is done by subtracting the measured intensity of a control Au sample with no protein from the overall protein sample fluorescence. While the fluorophore and imaging settings used for fluorescence images of both protein-functionalized samples were the same, the number of fluorophores per protein molecule, represented by the DOL, was different for each. As such, it was necessary to scale the subtracted intensities by the DOL for each protein to obtain the normalized intensity, or the intensity that would have been measured had each protein possessed exactly one fluorophore conjugate to it. Uncertainties in <I_protein, norm_> were calculated by calculating the standard errors for <I_protein_> and <I_Au Control_> from their respective standard deviations and following standard error propagation rules.

Bright pixels, such as those associated with aggregates, were identified and ignored in further analysis by converting the raw images to masks via Max Entropy thresholding of fluorescence images and dividing the resulting image by 255 to create a binary mask (see Figure 2S). Raw fluorescence images were then multiplied by this mask. The percentage surface area occupied by aggregates was calculated from the masked images, dividing the masked surface area by the total surface area. The average coefficient of variation <CV> for each sample and the control were calculated via

Equation S6

$$\left\langle CV \right\rangle=\frac{\left\langle SD \right\rangle}{\left\langle I \right\rangle}$$

where <SD> is the average standard deviation from each image of a sample and <I> is the average pixel intensity of each image of a sample. As above, uncertainties in <SD> were calculated following standard error propagation rules.

**Supplemental Figures**

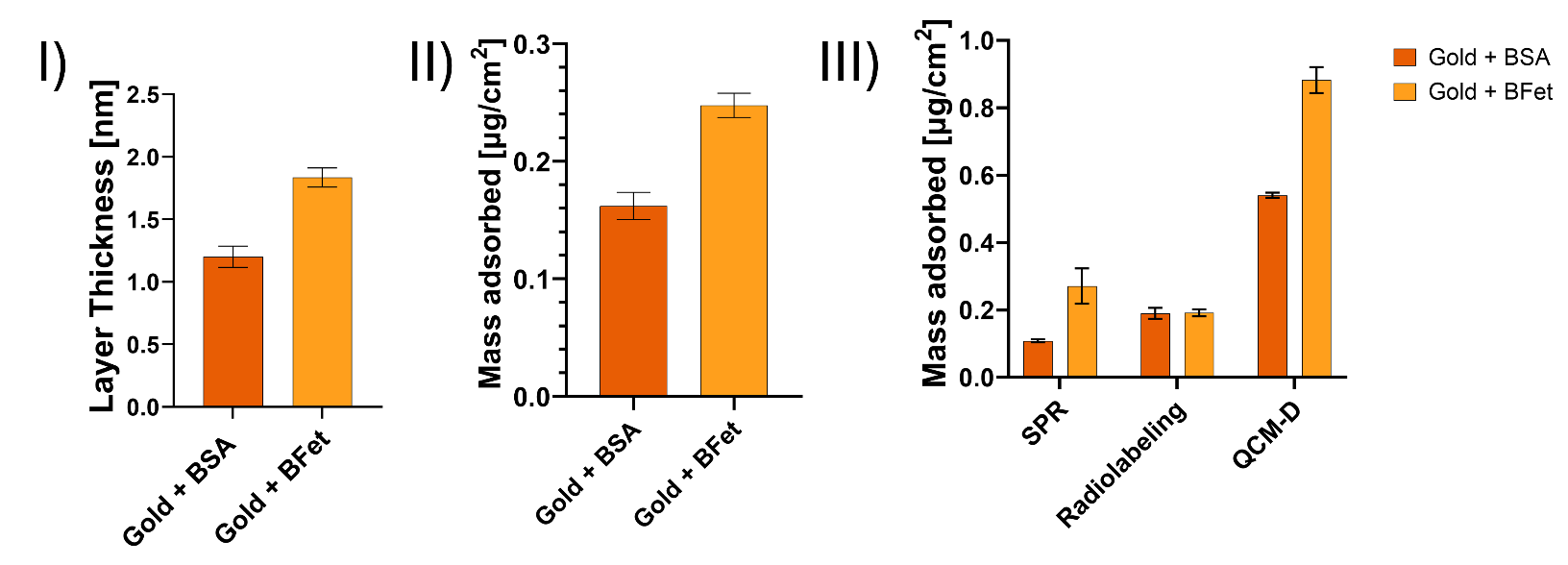

Figure 1S. Histograms represent I) the thickness of the protein layer evaluated from the change in SPR angle after 3 h submersion of the samples in 0.2 mg/mL protein solution, followed by PBS rinsing and nitrogen drying, II) the average protein mass adsorbed evaluated from the thickness data collected, assuming a 1.35 g/cm^3^ density of the protein layer. III) the average mass adsorbed for BSA (darker orange) and BFet (lighter orange) at plateau conditions for SPR and QCM-D (50 min injection of a 0.2 mg/mL protein solution in PBS) compared to the mass adsorption measured through radiolabeling (3 h adsorption from 0.2 mg/mL protein solution in PBS under static conditions).

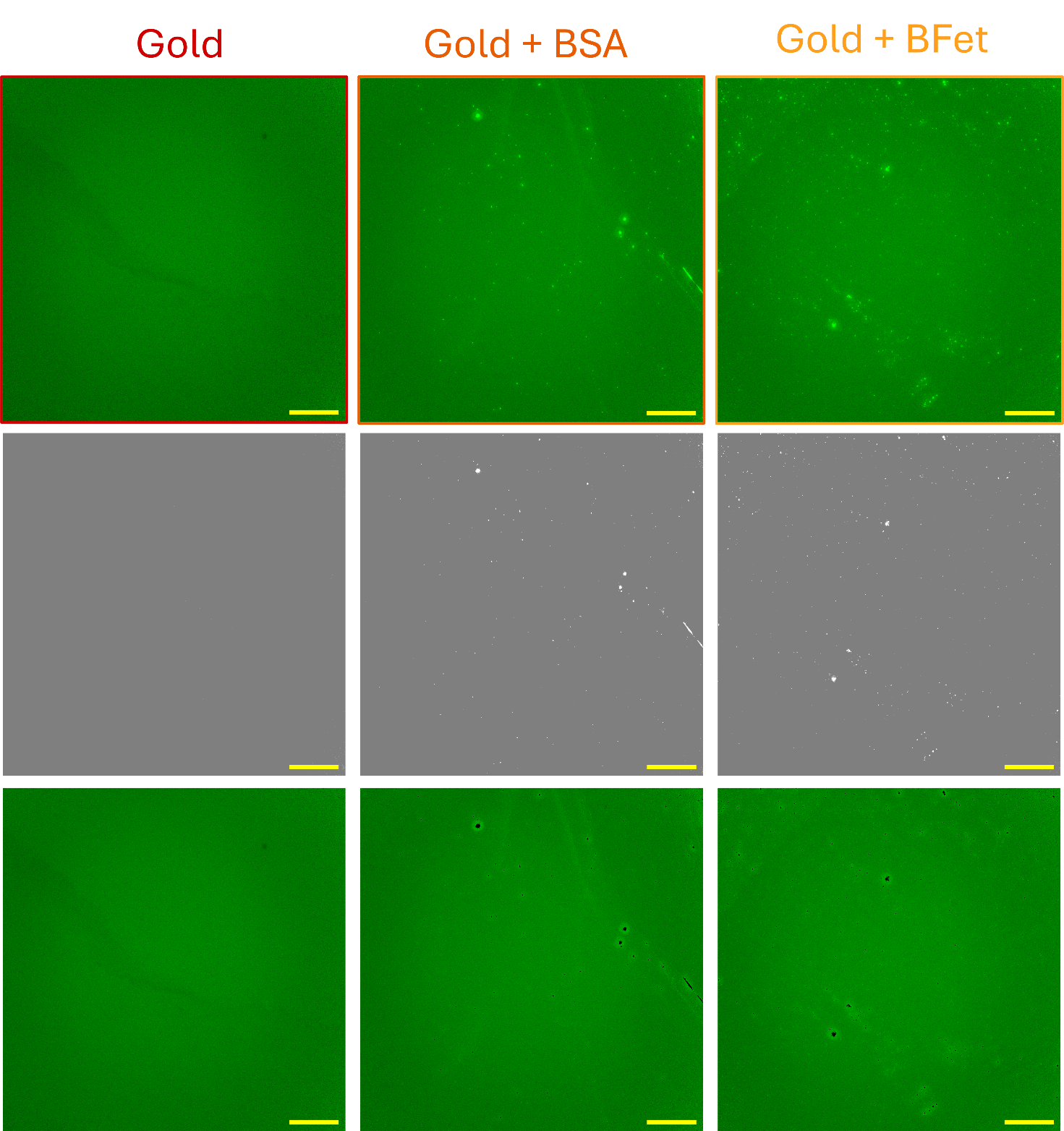

Figure 2S. The images present the workflow used for fluorescence intensity analysis for protein coverage distribution: the raw fluorescence micrographs (first row), max-entropy masks obtained with Fiji (second row) and final processed images (third row). The first column presents the resulting procedure for bare gold substrates, the second for Gold+BSA and the third for Gold+BFet.

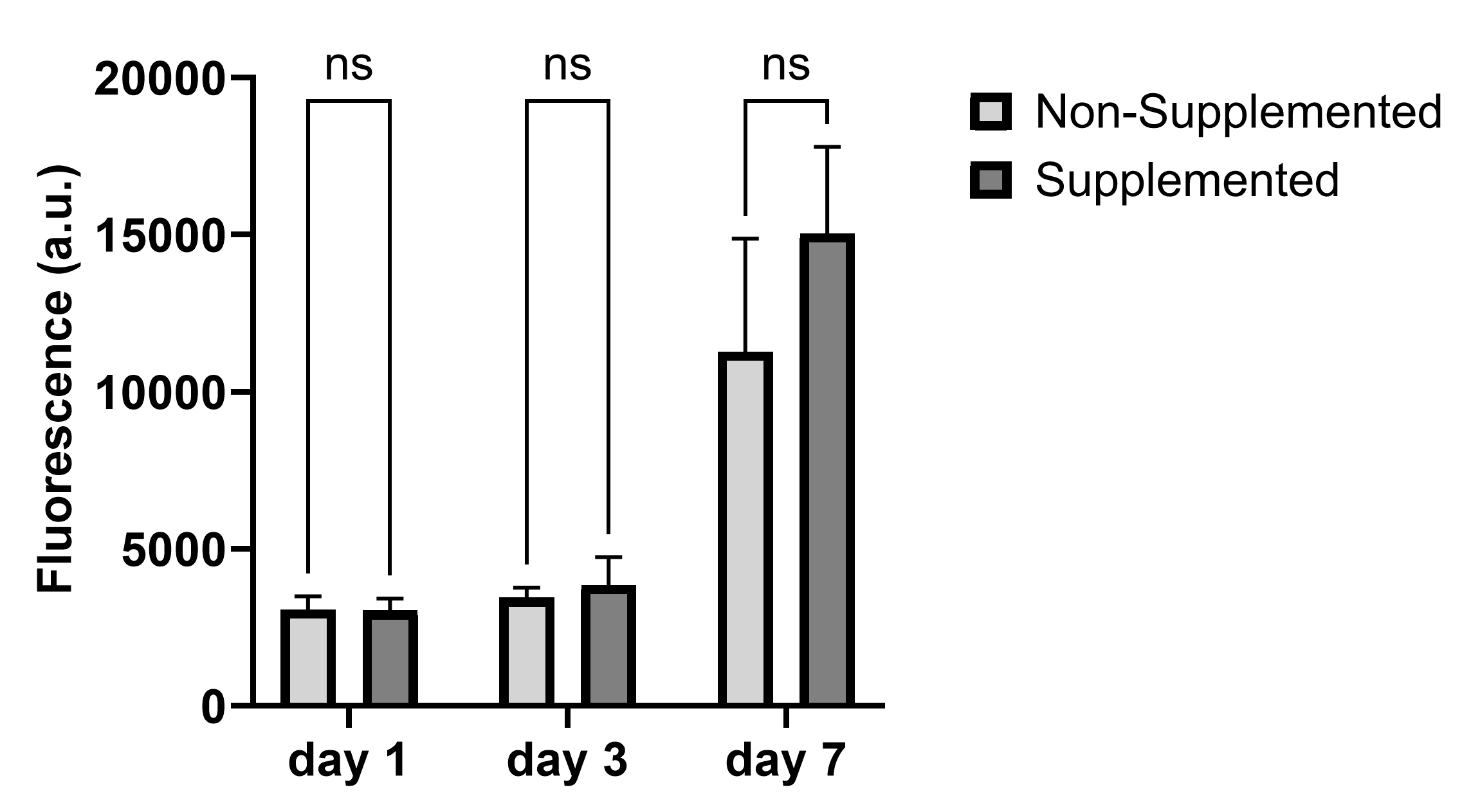

Figure 3S. The histogram describes the cellular metabolism measured through an Alamar Blue™ assay over 7 days for cells grown in 15%FBS-1%PS supplemented media (dark grey) or those cultured for 24 h in non-supplemented media and then in 15%FBS-1%PS supplemented media. The lack of significant differences in measured fluorescence over 7 days indicates no detrimental effect on the cells with the use of non-supplemented media for 24 h to avoid substitution of the pre-adsorbed protein.

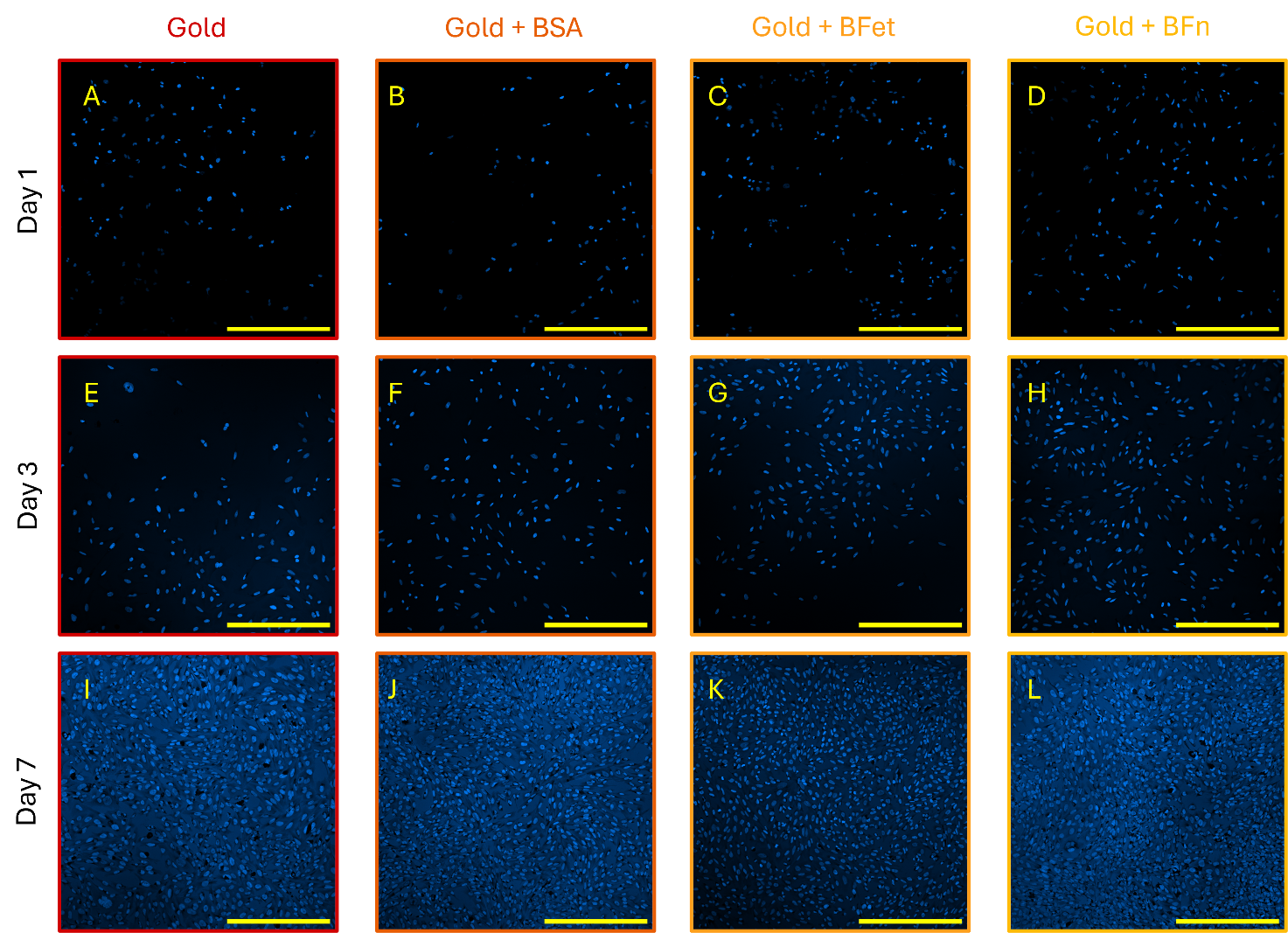

Figure 4S. These representative fluorescence micrographs show the number of cells present on substrates on day 1 (A-D), day 3 (E-H) and day 7 (I-L) from seeding. The first column (A-E-I) presents the cellular proliferation of Saos2 cells on bare gold surfaces, the second (B-F-J) proliferation on gold surfaces with pre-adsorbed BSA, the third (C-G-K) the proliferation on gold surfaces with pre-adsorbed BFet, and the last column (D-H-L) the proliferation of cells on gold surfaces with pre-adsorbed BFn. Scale bars correspond to 400 µm.
